# Power-to-power cross-frequency coupling as a novel approach for temporal lobe seizure detection and analysis

**DOI:** 10.1101/2025.05.31.657189

**Authors:** Bar Lehmann, Andrei V. Medvedev

## Abstract

**Objective:** Power-to-power cross-frequency coupling (CFC) is a novel method to index the dynamic spatio-temporal interactions between brain rhythms, including high frequency oscillations (HFOs). This research evaluates this promising method’s capacity for seizure detection with intracranial EEG. Seizures can be conceptualized as composites of different electrographic patterns including (1) spike, (2) ripple-on-spike, and (3) ripple-on-oscillation. This study also performs a basic CFC analysis of each of these components which has potential to further the understanding of epileptogenic processes.

**Methods:** In this study, deep learning networks including Stacked Sparse Autoencoder (SSAE) and Long Short Term Memory (LSTM) are trained to detect seizures and help characterize CFC patterns for these three common seizure components. The analysis uses intracranial EEG (iEEG) records from the ieeg.org (Mayo Clinic files) database. Temporal Lobe Epilepsy (TLE) seizures (*n* = 120) from 26 patients were analyzed along with segments of background activity. Power-to-power coupling was calculated between all frequencies 1-250 Hz pairwise using the EEGLAB toolbox. CFC matrices of seizure and background activity were used as training or testing inputs to the autoencoder.

**Results:** The trained network was able to recognize background and seizure segments (not used in training) with a sensitivity of 90.2%, specificity of 96.8% and overall accuracy of 93.4%. The three seizure components (spike, ripple-on-spike, ripple-on-oscillation) were also observed to have unique CFC signatures.

**Conclusions:** The results provide evidence both for (1) the relevance of power-to-power coupling (PPC) for TLE seizure detection in iEEG, as well as (2) there existing unique PPC signatures of three common seizure components.

## 1. Introduction

A wide range of frequencies from slow DC shifts (below 1Hz) to fast ripples (250-600Hz) characterize the human EEG (Park and Hong, 2019). Cross frequency coupling (CFC) refers to correlations between different brain oscillations such as between the phase of one oscillation and the amplitude of another (phase amplitude coupling), or the amplitude of one oscillation with the amplitude of another (power to power coupling) (Bernardo et al., 2020; Cai et al., 2021; Li et al., 2021). Such coordination between wavebands reflect a complex hierarchical organization of brain activity that spans not only the vast temporal scales mentioned (0-600Hz) but also the spatial scales from micro scale (individual neurons) to macro (brain networks) (Buzsaki and Draguhn, 2004; Klimesch, 2013). The dynamics of this cross frequency organization are likely to, at least partially, underpin both functional as well as pathological brain activity (Buzsaki and Draguhn, 2004).

Cross-frequency patterns offer a uniquely wide and useful net for brain research in that they can tie together multiple, and hitherto thought to be isolated, neurobiological features. In epilepsy for example, EEG slower oscillations that co-occur with another faster oscillation may reflect two separate neuronal populations without statistical or other functional interaction, or it may represent one neuronal population modulating another. CFC analysis can help us parse out which is correct. CFC analyses can capture a range of important characteristics of the coupling (amplitude, frequencies, phases) that together provide key clues into the meaning of the interaction such as which neuronal cell types, brain circuits, or networks are involved. It can also help us to understand the role of the coupling (such as propagation of activity from one brain region to another).

CFC interactions are in the juncture of many core questions about physiological brain function and also epilepsy. For example, since epilepsy is often thought of as a network disease (Sinha et al., 2022), and networks appear to operate in part if not primarily through CFC interactions (Besio et al., 2010; Bragin et al., 1999; Frauscher et al., 2017; Gardner et al., 2007; Luo et al., 2024; Medvedev et al., 2019; Siebenhuhner et al., 2020; Staba et al., 2014; Thomschewski et al., 2019; Worrell et al., 2004; Zijlmans et al., 2012), CFC analyses are poised to assist with understanding unknowns about how networks work or malfunction. While brain networks where discharges spread are often characterized as interconnected brain regions, this research is mostly based on BOLD signal correlations rather than cross frequency connection patterns, thus the CFC interactions that seem to underpin these networks remain to be uncovered. Another fundamental problem to understand is whether seizures in general (and their key features of spikes and higher frequency activity) can be characterized more as malfunctions of hyper-or hypoconnectivity. Formerly it was thought that a seizure was a disorder of increased connectivity/synchronization, however our group and others have probed different features and phases of seizures and found more nuance here. For example, we found indications that spikes may be an anti-synchronization mechanism, and other groups too have found increased desynchronization to be characterizing certain phases of a seizure (Medvedev, 2002; Wendling et al., 2003). While seizures may have much to do with increased synchrony locally, some research also suggests the seizure onset zone in particular is asynchronous/decoupled from the rest of brain activity (Wendling et al., 2003).

In the past decades, high frequency oscillations (HFOs), a waveband often defined between 30-600Hz (Engel and da Silva, 2012), have been revealed to have great significance for understanding cognitive activity (Medvedev and Kanwal, 2008; Pail et al., 2020). Increase in HFOs have also been found in pathologies, and the relationship between them and epilepsy has now been well documented (Blanco et al., 2010; Medvedev, 2002; Medvedev et al., 2011). For example, HFOs appear to increase before a seizure begins and can indicate the severity of the seizure disorder (with increased HFOs predicting more severe epilepsy) (Bernardo et al., 2020). While HFOs are not always routinely evaluated, removing HFO generating brain areas in patients with intractable localized epilepsy is correlated with better surgical outcomes than traditional methods (i.e. evaluating spikes) (Besio et al., 2010; Bragin et al., 1999; Medvedev et al., 2019; Thomschewski et al., 2019).

Cross frequency analyses between HFO and other waves (e.g. spikes or slower oscillations) have further refined our understanding in the area of epilepsy research. From enhancing automated detection of seizure vs. background activity, localization of the seizure onset zone (SOZ), prediction of seizure severity, differentiating between pathological vs. non-pathological HFOs, emerging research shows such CFC analyses appear to be a lens with much potential (Amiri et al., 2019; Iimura et al., 2018; Nariai et al., 2011). Mapping out the complex coupling relationships between the HFO and other wavebands in epilepsy provides important clues for understanding connectivity interactions of seizures. Such interactions may underlie core unanswered questions about epilepsy, such as how discharges propagate from the onset zone to the rest of the brain (Kohling and Staley, 2011; Matarrese et al., 2023; Muller et al., 2014).

The significance of using CFC with HFOs in epilepsy lies also in its ability to index various interrelated micro-to-macro scale processes (Li et al., 2022). The frequency metric of CFC can index neuronal cell types and mechanisms (Kawaguchi, 2001), while the coupling aspect can be localized to gauge what brain regions may be connecting. Formerly it was thought that, as opposed to slower frequencies that can synchronize brain regions across large spatial scales, the faster HFO frequencies represented a micro-scale and local-circuit agitation (Li et al., 2022). It now appears that large scale physiological synchronization by these HFOs is physiological (Arnulfo et al., 2020). Presently certain HFOs including fast ripples (200Hz-600Hz) are thought to be generated by network mechanisms, more specifically by the synchronized action potential activity of smaller populations of neurons whose oscillations are at different phases (Kohling and Staley, 2011). This mechanism is in contrast to most other slower oscillations which are generated by extracellular post synaptic potentials of large groups of pyramidal cells firing in synchrony. CFC biomarkers that include HFO may be proxies not only for the onset zone(s), but also for this long range synchronization (Li et al., 2022) that may be a mechanism of seizure propagation.

Methods utilizing cross-frequency coupling are increasingly researched both for seizure detection and also to help comprehend the propagation of epileptic discharges in the brain (Amiri et al., 2019; Arnulfo et al., 2020; Bernardo et al., 2020; Cai et al., 2021; Iimura et al., 2018; Jacobs et al., 2018; Li et al., 2021; Nariai et al., 2011). The different types of coupling (i.e. power-to-power, power-to-phase, phase-to-phase, etc.) are thought to have independent neural mechanisms as well as different or complimentary functional significance (Jirsa and Muller, 2013). While many forms of CFC have not been well-researched, the best studied form of CFC is phase-to-amplitude coupling (PAC), which is well known to have an association with various cognitive processes related to memory and perception, as well as epilepsy (Edakawa et al., 2016; Li, 2016; Medvedev and Kanwal, 2008). While not pathological on its own, PAC between gamma (20-50Hz) and the lower frequency bands including theta and delta (1-7Hz) have been found to be associated with epilepsy (Fujita et al., 2022; Jacobs et al., 2018).

Power-to-power coupling (PPC) is another type of cross-frequency coupling having a solid research base (Ferraris et al., 2018; Sheremet et al., 2019; Wang et al., 2018) attesting to its significance, yet in contrast with PAC, there is a surprising lack of research in the area of epilepsy. PPC calculation is based on simple time-course correlations which lends itself well for real-time seizure detection. PPC is not only more computationally efficient than other methods but is less likely to be impacted by signal noise. This is because of its reliance on power rather than phase, the latter of which is more vulnerable to noisy signals (Giehl et al., 2021). Thus in light of its suitability for real-time processing, as well as to help bridge the gap in the literature in the area of PPC and epilepsy, we investigate PPC for seizure detection.

It is important to recognize that a seizure is a dynamic process or perhaps even an amalgam of events rather than a singular monolithic event. For this reason, we have also singled out the common epileptiform elements of (1) spike, (2) ripple-on-spike (RonS), and (3) ripple-on-oscillation (RonO), each of which could be mined for comprehensive CFC relationships. There is much that has been uncovered about these elements but large questions also remain. For example, in the past, spikes have been related to broad cortical areas and deep brain structures, and the initial slow phase of a spike has been thought of as potentially being a “generator” or a recruitment mechanism for larger neuronal populations into the seizure (Matarrese et al., 2023). Spikes found to have coherence with slower frequencies (2-50Hz) have been proposed as biomarkers for the neuronal recruitment process that propagates local disfunction into a seizure (Ibrahim et al., 2014). Alternatively, our group has been examining grounds suggesting that these spikes, instead of propagating synchrony, may actually be acting to counter unchecked high frequency hypersynchronization between neuronal populations (Medvedev, 2002). Ripples are a class of HFO thought to be more related to highly local microcircuits, and they have been found to sometimes involve connectivity between different HFO-hubs (Li et al., 2021). Ripple-on-oscillation are known to be involved in epileptogensis (in particular, oscillations including the slower 1-12Hz) (Cai et al., 2021; Ibrahim et al., 2014; Li et al., 2022). Thus ripples and ripple-on-oscillations as classes also allow us consider the CFC signals in the context of what we know of the physiology of relevant microcircuits and network interactions. Evaluating the PPC relationships of these elements, in addition to being leveraged for seizure identification, can also provide new insight onto these areas.

## 2. Methods

We used intracranial EEG (iEEG) data from 26 patients with temporal lobe epilepsy (TLE) recorded specifically for clinical purposes during their evaluation for surgery. The datasets were obtained from the open source database ieeg.org (Mayo Clinic files) and contain de-identified segments of iEEG from epilepsy patients of different age, sex, and ethnicity. The Institutional Review Board of Georgetown-MedStar approved this study and verified that it was done in accord with the needed regulations, laws, and ethical guidelines. The privacy rights of human subjects have been observed, and only de-identifiable data from the public database was used. All patients had TLE with focal complex or partial seizures which in some patients could evolve into secondary generalized tonic-clonic seizures. The demographic data for patients and some statistical data about the seizures selected for the analysis are presented in Table 1. The seizures in this database are annotated by neurologists including their start and stop times. In total, 120 seizures were taken into analysis along with segments of seizure-free background activity. Background segments were selected so that they matched the seizure segments both in number and duration for each patient. The average duration of the selected seizures (and the corresponding background segments) varied from 17 to 177 seconds. The number of electrodes and their locations were chosen by medical staff for evaluating the subject-specific clinically relevant brain areas that may be contributing to that individual’s seizures. The number of channels varied from 16–128 across patients. These channels often spanned the temporal, frontal, and parietal cortices.

**Table 1.** Demographic information and clinical features of patients’ TLE seizures

| *No.* | *Sex* | *Hand* | *Age at admission (years)* | *Age at onset of seizures (years)* | *Seizure Type* | *No. of seizures analyzed* | *Mean length of segments analyzed (s)* | *No. of channels* |
| --- | --- | --- | --- | --- | --- | --- | --- | --- |
| *M1* | F | R | 27 | 14 | FBTCS | 2 | 51 | 54 |
| *M2* | M | R | 25 | 22 | FIAS | 2 | 48 | 56 |
| *M3* | F | R | 13 | 1 | FIAS | 2 | 130 | 54 |
| *MC4* | F | R | 34 | 10 | FIAS | 4 | 45 | 87 |
| *M5* | M | R | 33 | 23 | FIAS | 2 | 25 | 60 |
| *M6* | M | R | 37 | 23 | FIAS | 5 | 42 | 84 |
| *M7* | F | R | 33 | 9 | FIAS | 2 | 110 | 104 |
| *M8* | F | L | 36 | 5 | FBTCS | 6 | 35 | 64 |
| *M9* | M | R | 39 | 29 | FBTCS | 7 | 101 | 16 |
| *M10* | M | R | 33 | 31 | FBTCS | 9 | 135 | 96 |
| *M11* | M | R | 10 | 5 | FBTCS | 8 | 129 | 56 |
| *M12* | M | R | 16 | 13 | FBTCS | 4 | 48 | 108 |
| M13 | F | L | 21 | 19 | FBTCS | 6 | 43 | 55 |
| M14 | M | R | 16 | 1 | FIAS | 2 | 63 | 88 |
| M15 | F | R | 23 | 9 | FIAS | 15 | 41 | 88 |
| M16 | M | ambidextrous | 9 | 9 | FIAS | 1 | 33 | 96 |
| M17 | F | R | 34 | 21 | FBTCS | 3 | 91 | 48 |
| M18 | M | L | 5 | 4 | FBTCS | 2 | 21 | 96 |
| M19 | F | R | 22 | 19 | FBTCS | 3 | 89 | 64 |
| M20 | F | R | 18 | 16 | FBTCS | 7 | 71 | 64 |
| M21 | M | R | 5 | 5 | FBTCS | 2 | 46 | 116 |
| M22 | M | ambidextrous | 3 | 0 | Generalized/<br>Atonic | 8 | 177 | 128 |
| M23 | M | R | 45 | 25 | FBTCS | 3 | 135 | 92 |

FIAS, focal impaired awareness seizures; FBTCS, focal to bilateral tonic-clonic seizures

Only very problematic segments having big motion-related artifacts (signal shifts) and bad channels due to technical problems were excluded. Apart from this, the iEEG segments were taken into analysis as raw records without any preprocessing or filtering of any kind. Avoiding pre-processing steps in iEEG is feasible since the intracranial data is cleaner than scalp EEG, and also avoiding extra processing can help maximize computational efficiency for algorithms that may eventually be run in real time. In the majority of patients, the sampling frequency was 500 Hz. However, for a few where it was higher, the data were downsampled to 500 Hz in order to make the analysis uniform. The analysis was performed using a modified script based on the PowPowCAT toolbox from EEGLAB (Thammasan and Miyakoshi, 2020). The spectrogram for each EEG segment was computed using the short-time Fourier transform via the Matlab spectrogram function. This was performed with one-second epochs that overlapped by 50%, employing a Hamming window and covering frequencies ranging from 1 to 250 Hz on a logarithmic scale. The resulting spectrogram represented the temporal modulations of spectral power across each frequency, EEG channel, and segment. To quantify power-to-power coupling, Pearson correlation coefficients were computed between the spectral power time courses of all frequency pairs within each EEG channel, yielding channel-specific PPC matrices. These matrices were subsequently averaged across all iEEG channels, producing a PPC matrix for each iEEG segment. The segment-specific PPC matrices (of size 100×100) were converted into the 4950-point vectors by taking only the elements below the main diagonal (because PPC matrices are symmetrical around the main diagonal). These vectors were then used as training and testing sets for the detection of seizure versus background segments.

We used a Stacked Sparse Autoencoder (SSAE) deep learning network for seizure detection. This algorithm begins in an unsupervised way to find a lower-dimensional feature-set characterizing seizure vs. background state. This dimensionality reduction for these CFC input matrices may improve both computational efficiency and also detection by reducing overfitting to numerous non-essential features that may be particular to a subject, seizure-type, or sensor location. The SSAE was constructed using Matlab (v. R2023b) with two hidden encoder-decoder layers, and a softmax layer was used to detect each iEEG segment into the binary outputs of (1) ‘background’ or (2) ‘seizure.’ In each detection run, half of the patients (thirteen) were used for training and the other half was used for testing, and the selection of patients into each group was random. Ten classification runs were performed for cross-validation and the results were averaged across runs and then subjects to obtain the resultant confusion matrix with the final values of sensitivity, specificity, and accuracy (mean ± standard deviation). To further evaluate the performance of the SSAE detector, the receiver operating characteristic (ROC) curve was calculated using Matlab function *rocmetrics*.

An important question is how various distinctive epileptiform events in iEEG such as spikes, ripple-on-spike, and ripple-on-oscillation are reflected in the patterns of cross-frequency coupling. To calculate PPC matrices for those events separately, we applied the approach used in our previous study which is based on a Long Short Term Memory (LSTM) network ((Medvedev et al., 2019). First, the LSTM network was used in this study to detect spikes, RonS and RonO events. The events, separated from other events by at least 3.5 seconds in time, were selected and the PPC matrices were calculated on the 3-second non-overlapping epochs centered on each event. Finally,

the PPC matrices were averaged in each category of events for qualitative comparison. For example, such comparison helps answer the question of whether high frequencies, which are usually detected during epileptiform events, might be simply harmonics of the lower frequencies or be indicative of actual HFO bursts.

## 3. Results

Among the 26 patients included in this study, 15 were male and 11 were female. Patient ages ranged from 3 to 62 years, with a mean age of 26.5 ± 15.2 years. Demographic information and clinical characteristics related to seizures are summarized in Table 1. Ten patients experienced focal impaired awareness seizures (FIAS), characterized by focal onset—typically involving the temporal and/or frontal electrodes—with rapid secondary generalization. Fifteen patients presented with focal to bilateral tonic-clonic seizures (FBTCS), and one patient had generalized or atonic seizures, as detailed in Table 1.

A typical iEEG trace during seizure, along with the iEEG spectrogram, is shown in Fig 1. Temporal lobe seizures quite often started with bursts of high frequency activity in the high gamma-ripple range (red oval in Fig 1; seizure onset marked by the vertical dashed red line), and this was followed by rhythmic bursts with different and complex spectral patterns containing various frequencies (black oval in Fig 1). As the seizure progressed, these patterns tended to evolve spatially and temporally, often spreading to adjacent cortical regions and exhibiting waxing and waning amplitudes. In several cases, the initial high-frequency discharges gradually transitioned into lower frequency oscillations, including theta and delta components, particularly during the later phases of the seizure. These shifts in frequency were frequently accompanied by increased synchronization across recording channels, suggesting broader network involvement. Notably, the variability in spectral evolution across seizures and subjects underscores the heterogeneity of seizure dynamics, which may reflect differences in underlying pathophysiology or cortical excitability (Fig 1).

**Figure 1.**
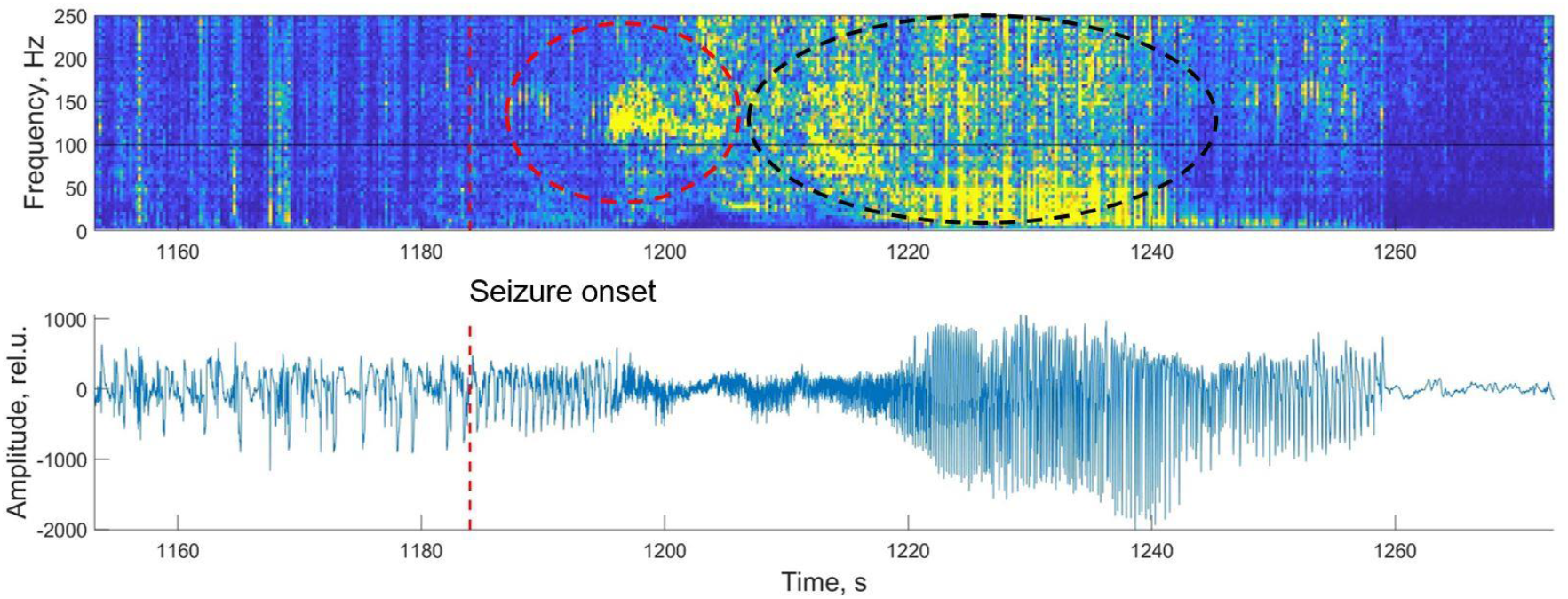
(*Bottom*) iEEG trace of seizure of one patient. (*Top*) spectrogram of this iEEG trace. Seizure onset is marked by vertical red line. Two ovals in the top and their corresponding trace below show how markedly heterogeneous portions of a single seizure can be.

A representative example of the coupling dynamics observed during seizures is shown in Fig 2, which illustrates a power-to-power frequency coupling matrix. This matrix captures the interactions between different frequency bands and reveals a complex, structured pattern of spectral coupling. Notably, all three major frequency bands commonly associated with seizure activity are evident: (a) the low-frequency band (3–13 Hz), encompassing delta to alpha rhythms (the black oval); (b) the beta to low-gamma band (the brown oval); and (c) the high-gamma to ripple band (the red oval), which includes fast oscillations often linked to epileptogenic activity. The matrix also highlights the presence of both the fundamental and second harmonic of a slow oscillation around 5 Hz (marked by the thick black arrows), which may reflect rhythmic spike discharges and possible harmonic coupling mechanisms (Fig 2).

**Figure 2.**
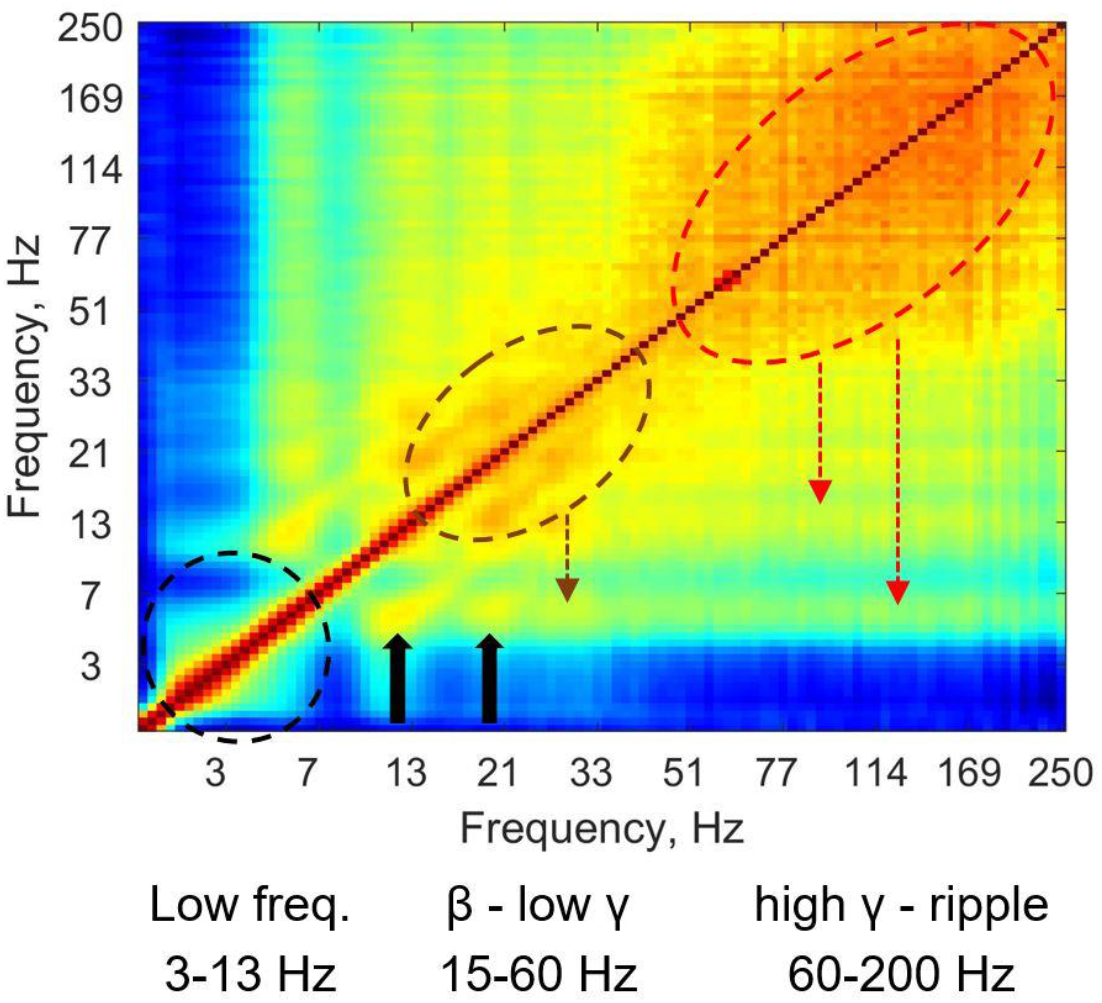
Cross-frequency matrix of iEEG activity during a TLE seizure. Circles, ovals and arrows show examples of relatively stronger coupling between different ranges of frequencies. 1^st^ and 2^nd^ harmonics of the slow rhythm around 5Hz (thick black arrows) may represent repetitive spikes and harmonic coupling. Two other higher frequency clusters (brown and red ovals) likely represent real gamma bursts and ripples. Importantly, these clusters show coupling with theta frequencies, which may be real CFC.

Importantly, two distinct high-frequency clusters (b) and (c) are unlikely to be the harmonics of the slower activity and may correspond to genuine gamma bursts and ripple-band activity. Nevertheless, these clusters display consistent coupling with theta-range frequencies (approximately 4–8 Hz; marked by the brown and red lines; Fig 2), suggesting a potential cross-frequency interaction between theta and faster gamma-ripple oscillations.

To uncover the patterns of power-to-power coupling associated with three major types of epileptiform events—spike, ripples-on-spike, and ripples-on-oscillations—we analyzed 8,900 segments of iEEG activity across all patients, seizures, and channels using an LSTM neural network (Medvedev, 2019). These events occurred with approximately equal frequency across the dataset. Figure 3 presents the corresponding averaged CFC matrices. The matrix for spike events (Fig. 3A) revealed a prominent cluster of coupling within the beta to low gamma frequency range (7–50 Hz; yellow oval in Fig 2A). Notably, this cluster may reflect harmonic coupling, a phenomenon arising due to the non-sinusoidal nature of repetitive spike waveforms. Because such waveforms contain harmonics that are strictly time-locked to the fundamental rhythm, they can produce significant – but spurious – coupling, which does not necessarily indicate interaction between distinct neural oscillations. Similar coupling patterns were observed in waveforms containing ripples (RonS and RonO; Fig 3, B–C, yellow ovals). However, both ripple-on-spike and ripple-on-oscillation events additionally exhibited pronounced coupling in the high gamma to ripple frequency range (60–250 Hz) (Fig. 3B–C, red ovals), suggesting the presence of more functionally relevant interactions in these higher frequency bands.

**Figure 3.**
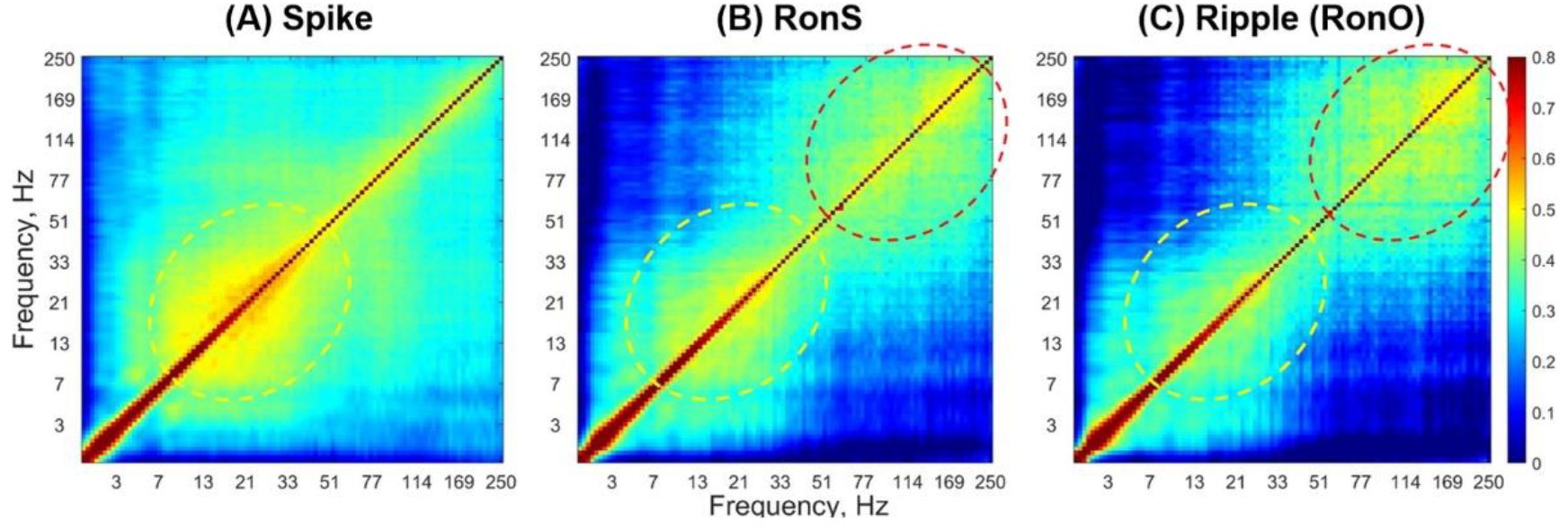
Cross-frequency coupling matrices characterizing the three seizure components. Power-to-power matrices are group-averaged over all instances of spike **(A)**, ripple-on-spike **(B)**, and ripple-on-oscillation **(C)** respectively. Yellow circles highlight lower frequency to low-gamma range, and red circles highlight the high-gamma to ripple range. These were all processed using a logarithmic frequency scale for optimal visualization.

The suggestion that the observed coupling patterns (Figs. 2–3) reflect epileptic waveforms is supported by a strong correlation between average coupling and the number of epileptiform events across three categories (spike, RonS and RonO). Using the LSTM network, we identified 3277 spikes, 2528 RonS events, and 3104 RonO events within the analyzed data segments. For each segment, we computed the event-specific average coupling as the mean value of the corresponding PPC matrix thus obtaining the segment-specific PPC metric. This PPC metric was then correlated with the number of events in each segment. The results indicate that cross-frequency coupling is strongly correlated with all three types of epileptic waveforms (Fig. 4). Correlation is strongest for spike and ripple-on-spike (Fig 4A, B) which is likely to be a result of the presence of harmonics of the sharp spike waveforms. For the ripple-on-oscillation waveform, the correlation is somewhat lower due to larger scattering of individual values. Nevertheless, the RonO events also show a highly significant correlation between the number of events and the corresponding PPC metric (Fig 4C). Significant cross-frequency coupling for the RonO events, where spurious harmonic coupling is less likely due to the lesser presence of sharp waveforms and a greater time jitter between high- and low-frequency waveforms, may indicate a physiologically meaningful coupling mechanism (not just the presence of harmonics). With this mechanism at work, slower oscillations modulate the amplitude or timing of faster rhythms—a phenomenon often implicated in the coordination of neuronal ensembles during seizure propagation. The structured appearance of these couplings across multiple seizures and subjects further supports their potential significance in the underlying seizure dynamics.

**Figure 4.**
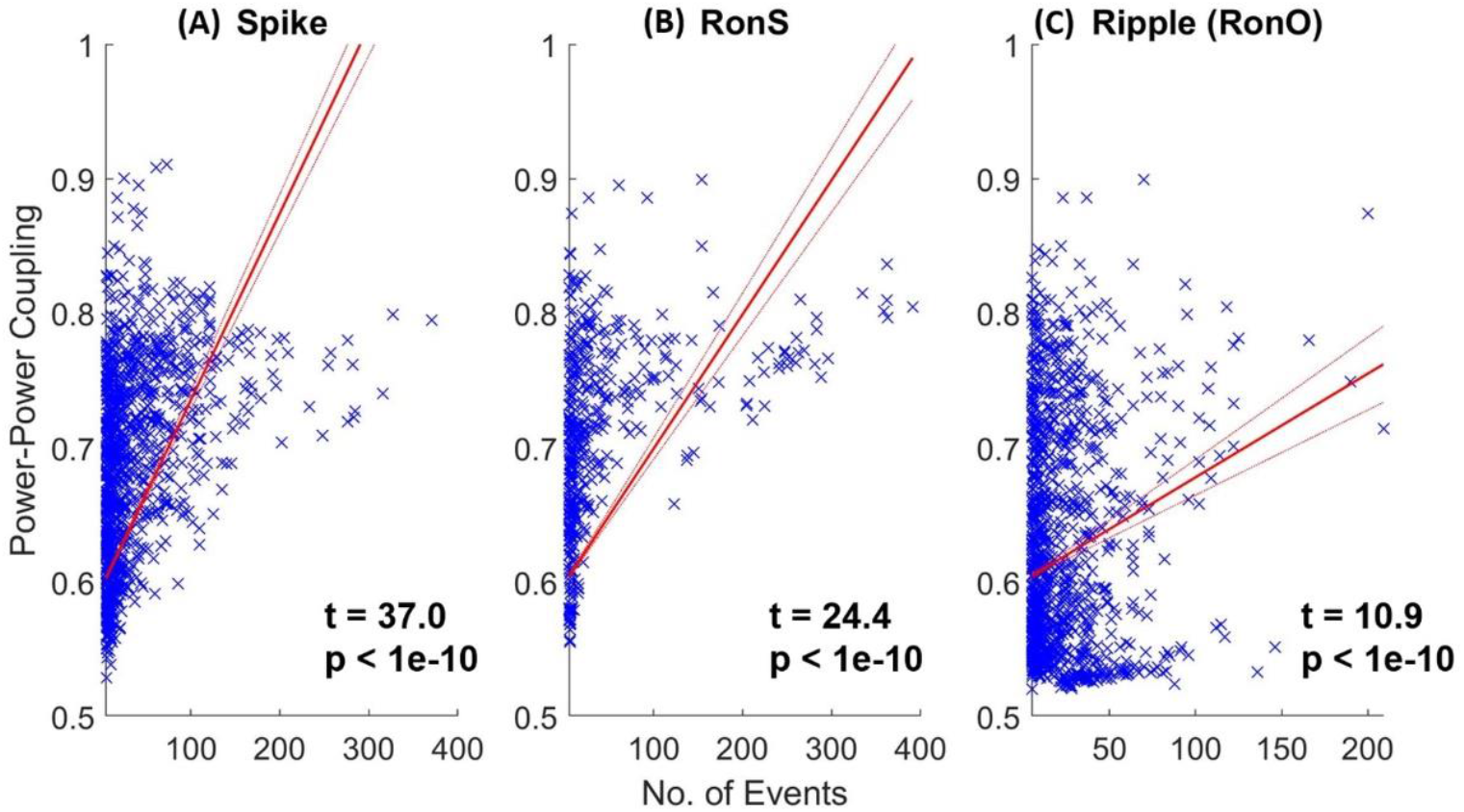
Correlation between average coupling and the number of events in each of the three categories: spike **(A)**, ripple-on-spike **(B)**, and ripple-on-oscillation **(C)**. Average coupling was calculated as a mean value of the segment-specific CFC matrix. This measure was correlated with the number of events in the corresponding segments. The high correlations observed highlight the stability of the relationship between these three seizure components their respective CFC patterns characterizing them across subjects, seizure types, and brain regions.

During training of the SSAE network with L2 and sparsity regularization, the squared error dropped below 10^−2^—the stopping criterion—after approximately 400 iterations. Fine-tuning was then performed. Figure 5 (*left*) presents the confusion matrix summarizing the SSAE network’s performance in detecting seizure versus background segments. On held-out iEEG data (not used in training), the trained network achieved an average sensitivity of 90.2%, specificity of 96.8%, and overall accuracy of 93.4%. Based on the total duration of analyzed iEEG segments, the 3.2% false positive rate (100% – specificity) corresponds to approximately seven false alarms per hour. The ROC curve is shown in Figure 5 (*right*), with an area under the curve (AUC) of 0.96 ± 0.06 (mean ± standard deviation) for the two-class average.

**Figure 5.**
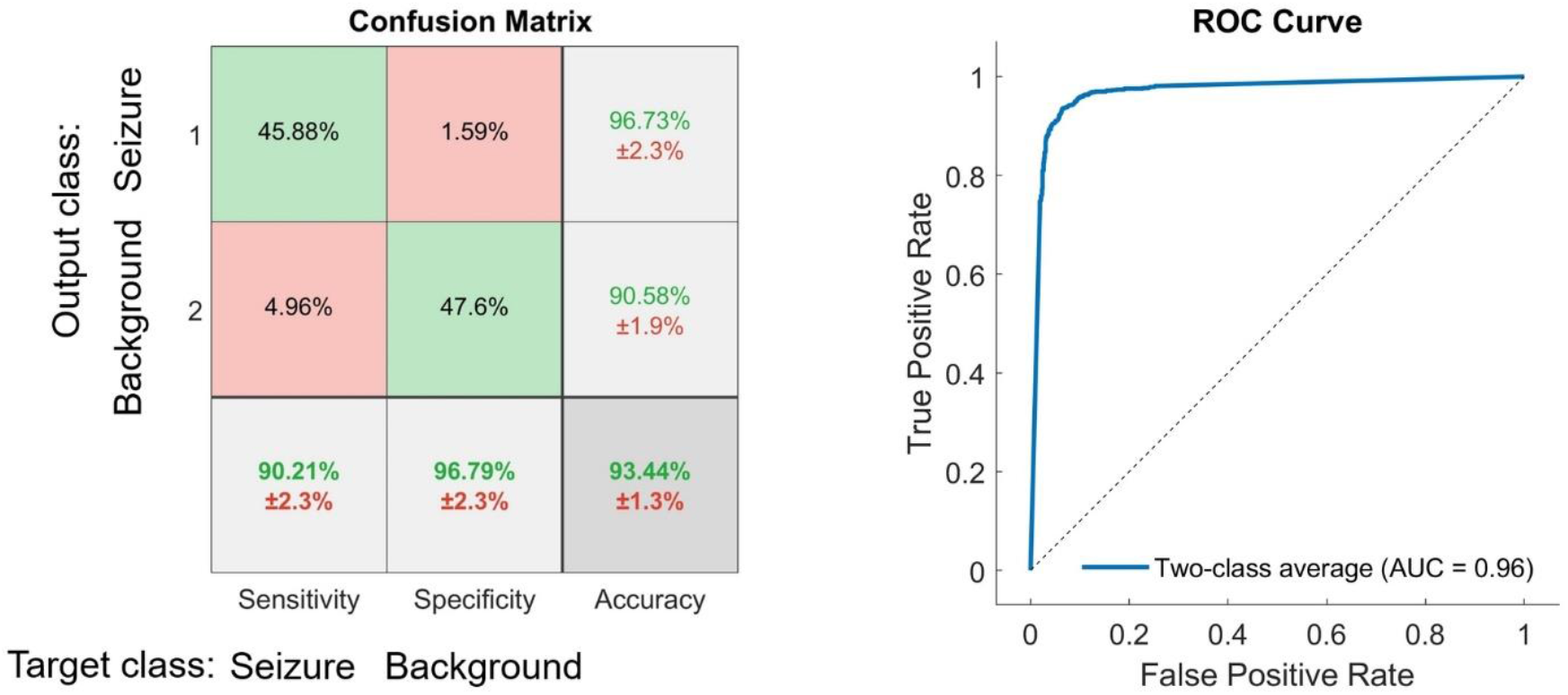
*Left:* Confusion matrix showing the results of recognition of seizures vs. background segments by the SSAE network. The mean values (%) ± standard deviations (%) are shown for sensitivity, specificity and overall accuracy (the bottom row) as well as for positive predictive values for each class (two upper cells in the right-hand column). *Right:* The ROC curve representing the classification results over both classes (seizure versus background) for the trained SSAE neural network. The area under the curve (AUC) metric is also shown.

## 4. Discussion

Detection of seizures is particularly important in TLE. TLE seizures may be electrographic yet lacking in obvious behavioral signs, and therefore can be easily missed by the medical staff (Medvedev et al., 2019). These patients are monitored for several days around the clock and the quantification of their seizures is a significant challenge. Despite the advanced tools of (iEEG) recordings, which have a higher fidelity and less noisy signals compared to magnetoencephalography (MEG) or scalp EEG, identifying seizures using visual inspection is very time-consuming. Such inspections may be prone to focus on large spikes but miss important biomarkers such as HFOs. This is because these occurrences can be particularly difficult to detect due to their smaller voltage as well as the very large datasets (recordings often span several-hours and up to 128-channels) (Medvedev et al., 2019).

For the task of automated seizure detection, many algorithms rely on description of the seizure EEG’s characteristic wavebands, amplitude changes (Kumar et al., 2021; Richard et al., 2015; Xanthopoulos et al., 2009), as well as harmonics (Abeysuriya et al., 2014; Lingli Hu, 2023); (Buteneers et al., 2013; Sitnikova et al., 2009). PPC is a method that has a high level of sensitivity to these features which are already known to have utility in seizure detection (i.e. specific frequency bands including HFOs, amplitudes changes, harmonics), and in addition, it provides important information on interrelationships between these features. Presently there are very few studies related to PPC in epilepsy. Our study helps uncover novel cross-frequency interactions within epileptiform activity. This has the potential to further refine sensitivity and specificity in seizure detection, especially in longer term monitoring where EEG is non-stationary.

Seizure detection algorithms including those using machine learning show increasingly positive results and have significant implications both for clinical application and etiological understanding. In an iEEG analyses using PAC with HFO’s Amiri et al. demonstrated accuracies around 85-90% using a support vector machine (SVM) algorithm (Amiri et al., 2019). Schirrmeister et al. detected seizures using spectral power between alpha and high gamma bands with a convolutional neural network (CNN) and achieved accuracies as high as 84% (Schirrmeister et al., 2017). Li et al. obtained an accuracy of 97.5% using multiscale entropy and Linear Discriminant Analysis algorithms (Li, 2016). Zhang et al. used a deep learning network which reached a sensitivity of 92.2% with a low false positives rate (FPR) of 0.12 per hour (Zhang et al., 2020). Other studies too have used various ML models for seizure prediction in human scalp EEG using different features with accuracy ranging from ∼70% to higher than 90-95% (Ansari et al., 2021; Liu et al., 2022; Sridevi et al., 2019; Thara et al., 2023; Yang et al., 2023).

Automated analysis is obviously of great value for real-time monitoring of epilepsy, and clearly machine learning techniques have made this appear more feasible. Deep learning (DL) stands out from other types of machine learning (ML) in that it is generally unsupervised, specialized for big datasets (including image matrices), complex features, and has superior ability to detect multifaceted latent patterns. For these reasons, it is not surprising that many successful detection studies have relied on various DL networks including Stacked Sparse Autoencoder (SSAE) (Liu et al., 2022; Schirrmeister et al., 2017; Zhang et al., 2020). This method is thus optimally suited for our aim of validating intricate CFC patterns for seizure detection. In our study, we used a SSAE which we trained to identify seizure (vs. non-seizure background) based on CFC patterns within iEEG. The sparseness of the SSAE refers to dimensionality reduction which is particularly useful when analyzing such large datasets.

We also utilized another DL algorithm, the Long Short Term Memory (LSTM) model, for collecting instances of seizure-related EEG patterns: (1) baseline, (2) spike, (3) ripple on spike, and (4) ripple on oscillation. The LSTM algorithm has been employed successfully for similar purposes of detecting brain time series data of cognitive states (Alhagry et al., 2017; Hasib, 2018). Furthermore, our group had demonstrated the value of this algorithm for detection of these particular categories in iEEG data with great generalizability across subjects and also datasets of two separate institutions (Medvedev et al., 2019). This time, after collecting these instances using this LSTM algorithm, we analyzed the cross frequency coupling signature of each of these.

Our method has built on the existing research in regard to (1) machine learning techniques, (2) the emerging understanding of HFO CFC and its relationship to epilepsy, and (3) using a method (PPC) that integrate various previously successful metrics (frequency bands, amplitudes, harmonics) in seizure detection. Our techniques’ accuracy of 93.44% is on par if not better than most other current detectors. By showing here our SSAE models’ accuracy in being able to detect TLE seizure vs. background activity with a hitherto unused metric (Power to Power CFC) we confirm that this metric has clear relevance to applications of epilepsy detection. Our inclusion of HFOs in our CFC analysis grounds our approach with recent epilepsy studies and the specific neurobiological significance of these faster waveforms that are described in those studies. This method appears to be robust to noise and has potential for clinical application in real-time monitoring (Giehl et al., 2021). We have also further validated two deep learning algorithms that have clear utility to such clinical application (SSAE and LMST).

This study highlights the value of an often overlooked metric (PPC) in epilepsy research, especially with the inclusion of HFO activity. In recent years there has been progress in mapping out the complimentary signatures of different modes of coupling (PAC, PPC, CFS, etc.) in cognition and pathology, which advances our understanding of brain dynamics (Arnulfo et al., 2020; Kawaguchi, 2001; Luo et al., 2024; Siebenhuhner et al., 2020). In this study we suggest additional PPC signatures to be included in this effort, specifically describing ripple-on-spike, ripple-on-oscillation, spike, and background activity. This outlines possible complimentary seizure biomarkers to the phase-amplitude-coupling patterns that are already known (Amiri et al., 2019; Iimura et al., 2018; Nariai et al., 2011).

There are a number of future directions that build on this research. There is reason to believe that PAC vs. CFS (cross frequency phase synchrony) are each related to distinct large scale brain networks each with different spectral features and separate hubs (Siebenhuhner et al., 2020). Mapping this complimentary coupling mode of PPC helps unravel the relationships between different frequency bands (including HFOs, which are already known to be deeply related to seizures), harmonic patterns, histological and network-level discharge propagation, and the complimentary (or redundant) roles of different modes of coupling (Abeysuriya et al., 2014; Arnulfo et al., 2020; Engel and da Silva, 2012; Iimura et al., 2018; Kawaguchi, 2001; Li et al., 2022; Lingli Hu, 2023; Luo et al., 2024; Nariai et al., 2011; Siebenhuhner et al., 2020; Sinha et al., 2022). For example, examining the PPC signatures of an antecedent spike vs. the subsequent between-spike period could help further test and elaborate on our previous research examining the possible role of spikes as a countering mechanism for pathological hyper-synchronization. This role of spikes was evaluated based on phase related metrics and could be better understood with such a complimentary lens of PPC (Medvedev, 2002). Further research in this vein may also contribute to more effective (and possibly less invasive) means to localize the SOZ for resection or neuromodulation.

The study limitations include first that all of the iEEG data is from a single institution (Mayo clinic). Second, while many of the PPC coupling patterns likely represent real coupling, testing the reality of coupling patterns, especially with non-sinusoidal morphologies and harmonics, requires more investigation. While harmonics are often unexamined or taken to be artifactual (which is certainly true some of the time), there is reason to think they can also contain salient information regarding neural dynamics of seizure propagation (Lingli Hu, 2023). Third, the physiological significance of the PPC in the context of epilepsy is another vital area that has not been well elucidated yet in our study and more generally. The coupling signatures may represent epiphenomenal byproducts of the seizure network, cellular mechanisms that inhibit the seizure, or mechanisms of seizure propagation / recruitment. For example, slower oscillations such as alpha frequency have known associations with cortical inhibition and so HFO coupling with this range may indicate a breakdown in inhibitory processes (Ibrahim et al., 2014). Fourth, we do not know how different this activity is from functional CFC patterns such as those normally occurring between HFOs and slower oscillations during memory-related processes (Ibrahim et al., 2014). Fifth, the physiological (and predictive) meaning of PPC signatures as redundant or complimentary to what is already known using PAC remains to be elucidated.

## Acknowledgments

The authors wish to thank the team of the iEEG.org data portal for giving us access to the data.

## Author contributions

BL: Data curation, Formal analysis, Investigation, Software, Writing – original draft, Writing – review & editing.

AM: Conceptualization, Data curation, Formal analysis, Funding acquisition, Investigation, Methodology, Project administration, Resources, Software, Supervision, Validation, Visualization, Writing – original draft, Writing – review & editing.

## Funding Sources

The author(s) declare that financial support was received for the research, authorship, and/or publication of this article. This work was supported by the National Institute of Mental Health of the National Institutes of Health under Grant Number RF1MH123192 to AM. The content is solely the responsibility of the authors and does not necessarily represent the official views of the National Institutes of Health.

## Generative AI statement

The authors declare that no Gen AI was used in the creation of this manuscript.

## Notes

https://www.ieeg.org

### Competing Interest Statement

The authors have declared no competing interest.

## References

Abeysuriya, R.G., Rennie, C.J., Robinson, P.A., Kim, J.W., 2014. Experimental observation of a theoretically predicted nonlinear sleep spindle harmonic in human EEG. Clin Neurophysiol 125, 2016–2023.

Alhagry, S., Fahmy, A.A., El-Khoribi, R.A., 2017. Emotion recognition based on EEG using LSTM recurrent neural network. Int J Adv Comput Sci Appl 8, 355–358

Amiri, M., Frauscher, B., Gotman, J., 2019. Interictal coupling of HFOs and slow oscillations predicts the seizure-onset pattern in mesiotemporal lobe epilepsy. Epilepsia 60, 1160–1170.

Ansari, A.Q., Sharma, P., Tripathi, M., 2021. A patient-independent classification system for onset detection of seizures. Biomed Tech (Berl) 66, 267–274.

Arnulfo, G., Wang, S.H., Myrov, V., Toselli, B., Hirvonen, J., Fato, M.M., Nobili, L., Cardinale, F., Rubino, A., Zhigalov, A., Palva, S., Palva, J.M., 2020. Long-range phase synchronization of high-frequency oscillations in human cortex. Nat Commun 11, 5363.

Bernardo, D., Nariai, H., Hussain, S.A., Sankar, R., Wu, J.Y., 2020. Interictal scalp fast ripple occurrence and high frequency oscillation slow wave coupling in epileptic spasms. Clin Neurophysiol 131, 1433–1443.

Besio, W., Gale, K., Medvedev, A., 2010. Possible therapeutic effects of transcutaneous electrical stimulation via concentric ring electrodes. 10th Workshop on Neurobiology of Epilepsy (WONOEP 2009). Epilepsia 51(S3), 85–87.

Blanco, J.A., Stead, M., Krieger, A., Viventi, J., Marsh, W.R., Lee, K.H., Worrell, G.A., Litt, B., 2010. Unsupervised classification of high-frequency oscillations in human neocortical epilepsy and control patients. J Neurophysiol 104, 2900–2912.

Bragin, A., Engel, J., Jr., Wilson, C.L., Fried, I., Mathern, G.W., 1999. Hippocampal and entorhinal cortex high-frequency oscillations (100--500 Hz) in human epileptic brain and in kainic acid--treated rats with chronic seizures. Epilepsia 40, 127–137.

Buteneers, P., Verstraeten, D., Nieuwenhuyse, B.V., Stroobandt, D., Raedt, R., Vonck, K., Boon, P., Schrauwen, B., 2013. Real-time detection of epileptic seizures in animal models using reservoir computing. Epilepsy Res 103, 124–134.

Buzsaki, G., Draguhn, A., 2004. Neuronal oscillations in cortical networks. Science 304, 1926–1929.

Cai, Z., Sohrabpour, A., Jiang, H., Ye, S., Joseph, B., Brinkmann, B.H., Worrell, G.A., He, B., 2021. Noninvasive high-frequency oscillations riding spikes delineates epileptogenic sources. Proc Natl Acad Sci U S A 118.

Edakawa, K., Yanagisawa, T., Kishima, H., Fukuma, R., Oshino, S., Khoo, H.M., Kobayashi, M., Tanaka, M., Yoshimine, T., 2016. Detection of Epileptic Seizures Using Phase-Amplitude Coupling in Intracranial Electroencephalography. Sci Rep 6, 25422.

Engel, J., Jr., da Silva, F.L., 2012. High-frequency oscillations - where we are and where we need to go. Prog Neurobiol 98, 316–318.

Ferraris, M., Ghestem, A., Vicente, A.F., Nallet-Khosrofian, L., Bernard, C., Quilichini, P.P., 2018. The Nucleus Reuniens Controls Long-Range Hippocampo-Prefrontal Gamma Synchronization during Slow Oscillations. J Neurosci 38, 3026–3038.

Frauscher, B., Bartolomei, F., Kobayashi, K., Cimbalnik, J., van ‘t Klooster, M.A., Rampp, S., Otsubo, H., Holler, Y., Wu, J.Y., Asano, E., Engel, J., Jr., Kahane, P., Jacobs, J., Gotman, J., 2017. High-frequency oscillations: The state of clinical research. Epilepsia 58, 1316–1329.

Fujita, Y., Yanagisawa, T., Fukuma, R., Ura, N., Oshino, S., Kishima, H., 2022. Abnormal phase-amplitude coupling characterizes the interictal state in epilepsy. J Neural Eng 19.

Gardner, A.B., Worrell, G.A., Marsh, E., Dlugos, D., Litt, B., 2007. Human and automated detection of high-frequency oscillations in clinical intracranial EEG recordings. Clin Neurophysiol 118, 1134–1143.

Giehl, J., Noury, N., Siegel, M., 2021. Dissociating harmonic and non-harmonic phase-amplitude coupling in the human brain. Neuroimage 227, 117648.

Hasib, M.M., Nayak, T. & Huang, Y., 2018. A hierarchical LSTM model with attention for modeling EEG non-stationarity for human decision prediction. IEEE EMBS International Conference on Biomedical & Health Informatics (BHI), 104–107

Ibrahim, G.M., Wong, S.M., Anderson, R.A., Singh-Cadieux, G., Akiyama, T., Ochi, A., Otsubo, H., Okanishi, T., Valiante, T.A., Donner, E., Rutka, J.T., Snead, O.C., 3rd, Doesburg, S.M., 2014. Dynamic modulation of epileptic high frequency oscillations by the phase of slower cortical rhythms. Exp Neurol 251, 30–38.

Iimura, Y., Jones, K., Takada, L., Shimizu, I., Koyama, M., Hattori, K., Okazawa, Y., Nonoda, Y., Asano, E., Akiyama, T., Go, C., Ochi, A., Snead, O.C., 3rd, Donner, E.J., Rutka, J.T., Drake, J.M., Otsubo, H., 2018. Strong coupling between slow oscillations and wide fast ripples in children with epileptic spasms: Investigation of modulation index and occurrence rate. Epilepsia 59, 544–554.

Jacobs, D., Hilton, T., Del Campo, M., Carlen, P.L., Bardakjian, B.L., 2018. Classification of Pre-Clinical Seizure States Using Scalp EEG Cross-Frequency Coupling Features. IEEE Trans Biomed Eng 65, 2440–2449.

Jirsa, V., Muller, V., 2013. Cross-frequency coupling in real and virtual brain networks. Front Comput Neurosci 7, 78.

Kawaguchi, Y., 2001. Distinct firing patterns of neuronal subtypes in cortical synchronized activities. J Neurosci 21, 7261–7272.

Klimesch, W., 2013. An algorithm for the EEG frequency architecture of consciousness and brain body coupling. Front Hum Neurosci 7, 766.

Kohling, R., Staley, K., 2011. Network mechanisms for fast ripple activity in epileptic tissue. Epilepsy Res 97, 318–323.

Kumar, A., Lyzhko, E., Hamid, L., Srivastav, A., Stephani, U., Japaridze, N., 2021. Differentiating ictal/subclinical spikes and waves in childhood absence epilepsy by spectral and network analyses: A pilot study. Clin Neurophysiol 132, 2222–2231.

Li, C., Sohrabpour, A., Jiang, H., He, B., 2021. High-Frequency Hubs of the Ictal Cross-Frequency Coupling Network Predict Surgical Outcome in Epilepsy Patients. IEEE Trans Neural Syst Rehabil Eng 29, 1290–1299.

Li, J., Liu, X. & Ouyang, G., 2016. Using Relevance Feedback to Distinguish the Changes in EEG During Different Absence Seizure Phases. Clin EEG Neurosci 47, 211–219.

Li, X., Zhang, H., Lai, H., Wang, J., Wang, W., Yang, X., 2022. High-Frequency Oscillations and Epileptogenic Network. Curr Neuropharmacol 20, 1687–1703.

Lingli Hu, L.Y., Hongyi Ye, Xiaochen Liu, Yuanming Zhang, Zhe Zheng, Hongjie Jiang, Cong Chen, Zhongjin Wang, Junming Zhu, Zhong Chen, Dongping Yang, Shuang Wang, 2023. Harmonic patterns embedding ictal EEG signals in focal epilepsy: a new insight into the epileptogenic zone. medRxiv 10.1101/2023.12.20.23300274

Liu, G., Tian, L., Zhou, W., 2022. Patient-Independent Seizure Detection Based on Channel-Perturbation Convolutional Neural Network and Bidirectional Long Short-Term Memory. Int J Neural Syst 32, 2150051.

Luo, Y., Meng, X., Zhou, G., Zhou, J., Luo, Y.J., Ai, H., Zelano, C., Chen, F., Xu, P., 2024. Oscillatory mechanisms of intrinsic human brain networks. Neuroimage 298, 120773.

Matarrese, M.A.G., Loppini, A., Fabbri, L., Tamilia, E., Perry, M.S., Madsen, J.R., Bolton, J., Stone, S.S.D., Pearl, P.L., Filippi, S., Papadelis, C., 2023. Spike propagation mapping reveals effective connectivity and predicts surgical outcome in epilepsy. Brain 146, 3898–3912.

Medvedev, A.V., 2002. Epileptiform spikes desynchronize and diminish fast (gamma) activity of the brain. An “anti-binding” mechanism? Brain Res Bull 58, 115–128.

Medvedev, A.V., Agoureeva, G.I., Murro, A.M., 2019. A Long Short-Term Memory neural network for the detection of epileptiform spikes and high frequency oscillations. Sci Rep 9, 19374.

Medvedev, A.V., Kanwal, J.S., 2008. Communication call-evoked gamma-band activity in the auditory cortex of awake bats is modified by complex acoustic features. Brain Res 1188, 76–86.

Medvedev, A.V., Murro, A.M., Meador, K.J., 2011. Abnormal interictal gamma activity may manifest a seizure onset zone in temporal lobe epilepsy. Int J Neural Syst 21, 103–114.

Muller, M.F., Rummel, C., Goodfellow, M., Schindler, K., 2014. Standing waves as an explanation for generic stationary correlation patterns in noninvasive EEG of focal onset seizures. Brain Connect 4, 131–144.

Nariai, H., Matsuzaki, N., Juhasz, C., Nagasawa, T., Sood, S., Chugani, H.T., Asano, E., 2011. Ictal high-frequency oscillations at 80-200 Hz coupled with delta phase in epileptic spasms. Epilepsia 52, e130–134.

Pail, M., Cimbalnik, J., Roman, R., Daniel, P., Shaw, D.J., Chrastina, J., Brazdil, M., 2020. High frequency oscillations in epileptic and non-epileptic human hippocampus during a cognitive task. Sci Rep 10, 18147.

Park, C.J., Hong, S.B., 2019. High Frequency Oscillations in Epilepsy: Detection Methods and Considerations in Clinical Application. J Epilepsy Res 9, 1–13.

Richard, C.D., Tanenbaum, A., Audit, B., Arneodo, A., Khalil, A., Frankel, W.N., 2015. SWDreader: a wavelet-based algorithm using spectral phase to characterize spike-wave morphological variation in genetic models of absence epilepsy. J Neurosci Methods 242, 127–140.

Schirrmeister, R.T., Springenberg, J.T., Fiederer, L.D.J., Glasstetter, M., Eggensperger, K., Tangermann, M., Hutter, F., Burgard, W., Ball, T., 2017. Deep learning with convolutional neural networks for EEG decoding and visualization. Hum Brain Mapp 38, 5391–5420.

Sheremet, A., Kennedy, J.P., Qin, Y., Zhou, Y., Lovett, S.D., Burke, S.N., Maurer, A.P., 2019. Theta-gamma cascades and running speed. J Neurophysiol 121, 444–458.

Siebenhuhner, F., Wang, S.H., Arnulfo, G., Lampinen, A., Nobili, L., Palva, J.M., Palva, S., 2020. Genuine cross-frequency coupling networks in human resting-state electrophysiological recordings. PLoS Biol 18, e3000685.

Sinha, N., Joshi, R.B., Sandhu, M.R.S., Netoff, T.I., Zaveri, H.P., Lehnertz, K., 2022. Perspectives on Understanding Aberrant Brain Networks in Epilepsy. Front Netw Physiol 2, 868092.

Sitnikova, E., Hramov, A.E., Koronovsky, A.A., van Luijtelaar, G., 2009. Sleep spindles and spike-wave discharges in EEG: Their generic features, similarities and distinctions disclosed with Fourier transform and continuous wavelet analysis. J Neurosci Methods 180, 304–316.

Sridevi, V., Ramasubba Reddy, M., Srinivasan, K., Radhakrishnan, K., Rathore, C., Nayak, D.S., 2019. Improved Patient-Independent System for Detection of Electrical Onset of Seizures. J Clin Neurophysiol 36, 14–24.

Staba, R.J., Stead, M., Worrell, G.A., 2014. Electrophysiological biomarkers of epilepsy. Neurotherapeutics 11, 334–346.

Thammasan, N., Miyakoshi, M., 2020. Cross-Frequency Power-Power Coupling Analysis: A Useful Cross-Frequency Measure to Classify ICA-Decomposed EEG. Sensors (Basel) 20, 7040.

Thara, D.K., Premasudha, B.G., Krivic, S., 2023. Detection of epileptic seizure events using pre-trained convolutional neural network, VGGNet and ResNet. Expert Systems e13447.

Thomschewski, A., Hincapie, A.S., Frauscher, B., 2019. Localization of the Epileptogenic Zone Using High Frequency Oscillations. Front Neurol 10, 94.

Wang, L., Hagoort, P., Jensen, O., 2018. Language Prediction Is Reflected by Coupling between Frontal Gamma and Posterior Alpha Oscillations. J Cogn Neurosci 30, 432–447.

Wendling, F., Bartolomei, F., Bellanger, J.J., Bourien, J., Chauvel, P., 2003. Epileptic fast intracerebral EEG activity: evidence for spatial decorrelation at seizure onset. Brain 126, 1449–1459.

Worrell, G.A., Parish, L., Cranstoun, S.D., Jonas, R., Baltuch, G., Litt, B., 2004. High-frequency oscillations and seizure generation in neocortical epilepsy. Brain 127, 1496–1506.

Xanthopoulos, P., Liu, C.C., Zhang, J., Miller, E.R., Nair, S.P., Uthman, B.M., Kelly, K., Pardalos, P.M., 2009. A robust spike and wave algorithm for detecting seizures in a genetic absence seizure model. Annu Int Conf IEEE Eng Med Biol Soc 2009, 2184–2187.

Yang, Y., Li, F., Qin, X., Wen, H., Lin, X., Huang, D., 2023. Feature separation and adversarial training for the patient-independent detection of epileptic seizures. Front Comput Neurosci 17, 1195334.

Zhang, Y., Guo, Y., Yang, P., Chen, W., Lo, B., 2020. Epilepsy Seizure Prediction on EEG Using Common Spatial Pattern and Convolutional Neural Network. IEEE J Biomed Health Inform 24, 465–474.

Zijlmans, M., Jiruska, P., Zelmann, R., Leijten, F.S., Jefferys, J.G., Gotman, J., 2012. High-frequency oscillations as a new biomarker in epilepsy. Ann Neurol 71, 169–178.

